# The Computational Grammar of Human Behaviour in a Biological Environment

**DOI:** 10.64898/2026.11.10.765265

**Authors:** Ulises Daniel Serratos Hernandez, Uzay Gökay, Lukas Kornemann, Yonatan Hutabarat, Debadrita Sen, Dominik R. Bach

## Abstract

Human behaviour unfolds as an extended stream of spontaneous actions shaped by environmental demands and the agent’s goals. The organisation of these actions, situated between setting abstract goals and concrete movement execution, remains poorly characterised. Here, we introduce DynaClade, an unsupervised temporal action segmentation algorithm for human full-body motion capture data, and apply it to freely moving participants navigating ecologically realistic risky foraging environments in virtual reality. DynaClade identifies discrete actions whose kinematic characteristics and spatial distributions correspond to environmental affordances, supporting the validity of the approach. Among these actions, we discovered a hitherto undescribed movement pattern defined by increased body sway without locomotion. Action sequences were highly non-random and showed clear history dependence beyond first-order Markov structure, most of which was explained by a second-order prediction. This higher-order structure was well captured in a hidden Markov model, in which some of the observed actions were associated with more than one latent state. Each of these latent states corresponded to the same action in a distinct action context. These findings suggest that spontaneous human actions are predominantly selected based on the current action and context, rather than following pre-planned or habitually assembled sequences.

## Introduction

Human behaviour unfolds in extended streams of movements, which enable individuals to pursue goals while dynamically adapting to their environments (Cooper and Shallice, 2006; Botvinick, 2008; Diedrichsen et al., 2026). From routine activities such as preparing a meal to complex social interactions, these movements are continuous, but temporally organized into identifiable actions. Despite theoretical work (Cooper and Shallice, 2006; Botvinick, 2008; Merel et al., 2019; Janssen et al., 2022) and empirical progress in understanding the generation of rehearsed and cued motor sequences (Helmich et al., 2009; Yokoi and Diedrichsen, 2019; Eckstein and Collins, 2021; Verwey, 2024; Diedrichsen et al., 2026), the organisation of spontaneous actions remains incompletely understood. While prospective planning of actions and movements is possible in structured laboratory tasks, it rapidly becomes intractable with extended state spaces and time horizons (Merel et al., 2019). Consistent with this, real-world observation of behavioural lapses in healthy and neurologically impaired persons has led to the suggestion that human actions are stochastically selected by model-free algorithms, shaped by contextual and goal demands rather than being pre-planned (Cooper and Shallice, 2000, 2006; Schwartz, 2006; Botvinick, 2008). This is consistent with limited planning depth even in fully cued or rehearsed sequences (Diedrichsen et al., 2026). As a related observation, spontaneous rodent behaviour can be modelled as a first-order Markov process, in which the selection of the next action depends only on the current one (Wiltschko et al., 2015), with action transition probabilities shaped by environmental conditions and internal states (Wiltschko et al., 2015; Weinreb et al., 2026). However, these non-human results may not directly generalise to human behaviour (Mobbs et al., 2021). At the same time, both in humans (Cooper and Shallice, 2000) and non-humans (Reddy et al., 2022), examples of fixed action sequences have been noted. In humans, these fixed sequences might arise by external task demands, by reflective planning (Mattar and Lengyel, 2022) converging on similar sequences in similar situations, or by a model-free controller that has access to the sequence of preceding actions. However, none of the aforementioned proposals have been tested on spontaneously generated human action sequences.

Understanding sequences of spontaneous actions calls for suitable experimental and data-analytic methods. This entails, first, observing and recording naturalistic action while retaining experimental control over context (Anderson and Perona, 2014; Krakauer et al., 2017). Second, it necessitates unsupervised methods to extract action sequences from continuous human movements in a situation where the action repertoire is not known a priori (Wiltschko et al., 2015; Hsu and Yttri, 2021; Ding et al., 2024). Of particular importance here are machine-learning methods, based on the key idea that although individual movements are rarely repeated precisely, actions can be grouped by spatiotemporal similarity (Ding et al., 2024; Gökay et al., 2025). In non-humans, these methods have dramatically enlarged the knowledge of behavioural structure and its underlying neural control (Markowitz et al., 2018, 2023; Geramita et al., 2026).

Here, we address this challenge using a computational ethology approach (Anderson and Perona, 2014). We record motion capture data while humans navigate ecologically realistic environments in freely-moving virtual reality (fm-VR), without imposing specific actions or predefined action orders. We develop DynaClade (**Dyna**mic hierarchical **Cl**ustering for **a**ction **d**iscovery in non-linear **e**mbeddings), an unsupervised temporal action segmentation algorithm that does not presuppose the action repertoire or the structure of action sequences. Importantly, DynaClade operates only on kinematic data with no information on the VR environment, ensuring that the assigned actions are not biased by what participants might be expected to do in a given context. We then analyse the resulting action repertoire and sequences.

A key ecological behaviour across species is foraging for food (Schoener, 1971; Hunt et al., 2021). Foraging is often shaped by external pressures, such as the presence of competing individuals or threatening predators (Mobbs and Kim, 2015; Bach et al., 2020; Bach, 2022; Trier et al., 2025). Risky foraging is well-suited for our purpose because it reliably elicits a rich and ecologically motivated repertoire of naturalistic actions (Sporrer et al., 2023). To capture the dynamics of risky foraging in an experimentally controlled manner, we expanded on previous work using fm-VR (Brookes et al., 2023; Sporrer et al., 2023; Hutabarat et al., 2025; Kornemann et al., 2026a) by tracking participants’ full-body movements while they navigated a wide variety of foraging environments containing potentially dangerous animals, weapons, and a safe shelter to which participants could escape. DynaClade extracts a meaningful repertoire of actions, allowing us to characterise the structure of action sequences in detail. We find robust evidence for higher-order sequence structure, but this is mainly explained by sensitivity to action context. Within an action context, actions appear to be selected largely according to a first-order Markov model.

## Results

### Detected actions align with task features

Participants navigated 94 distinct and self-contained fm-VR scenarios of 9-180 s (mean 35 s) duration (figure 1AB). We processed 1081 fm-VR epochs, corresponding to 10.6 hours of recordings and comprising overall 381,214 time bins at 10 Hz resolution from 23 participants. For each time bin, our unsupervised action segmentation algorithm, DynaClade, identifies three hierarchically nested action categories: a fine category distinguishing different movement execution styles, which are progressively collapsed at the intermediate and coarse level (figure 1C). To aid presentation and interpretation, we assigned descriptive labels to the action classes, based on inspection of representative exemplars of each identified category. Thus, DynaClade identified 22 coarse actions, 46 intermediate actions, and 70 fine actions (see examples in figure 1D and the full set of actions in figure 2). The average duration of an action bout was (mean ± standard deviation) 1.13 ± 2.67 s at the coarse level, 1.07 ± 2.50 s at the intermediate level, and 0.95 ± 2.00 s at the fine level, suggesting that these abstraction levels differ in spatial but less so in temporal granularity. In the following, unless noted otherwise, we focus on the coarse level, which had sufficient occurrences of every action for the analyses reported here.

**Figure 1:**
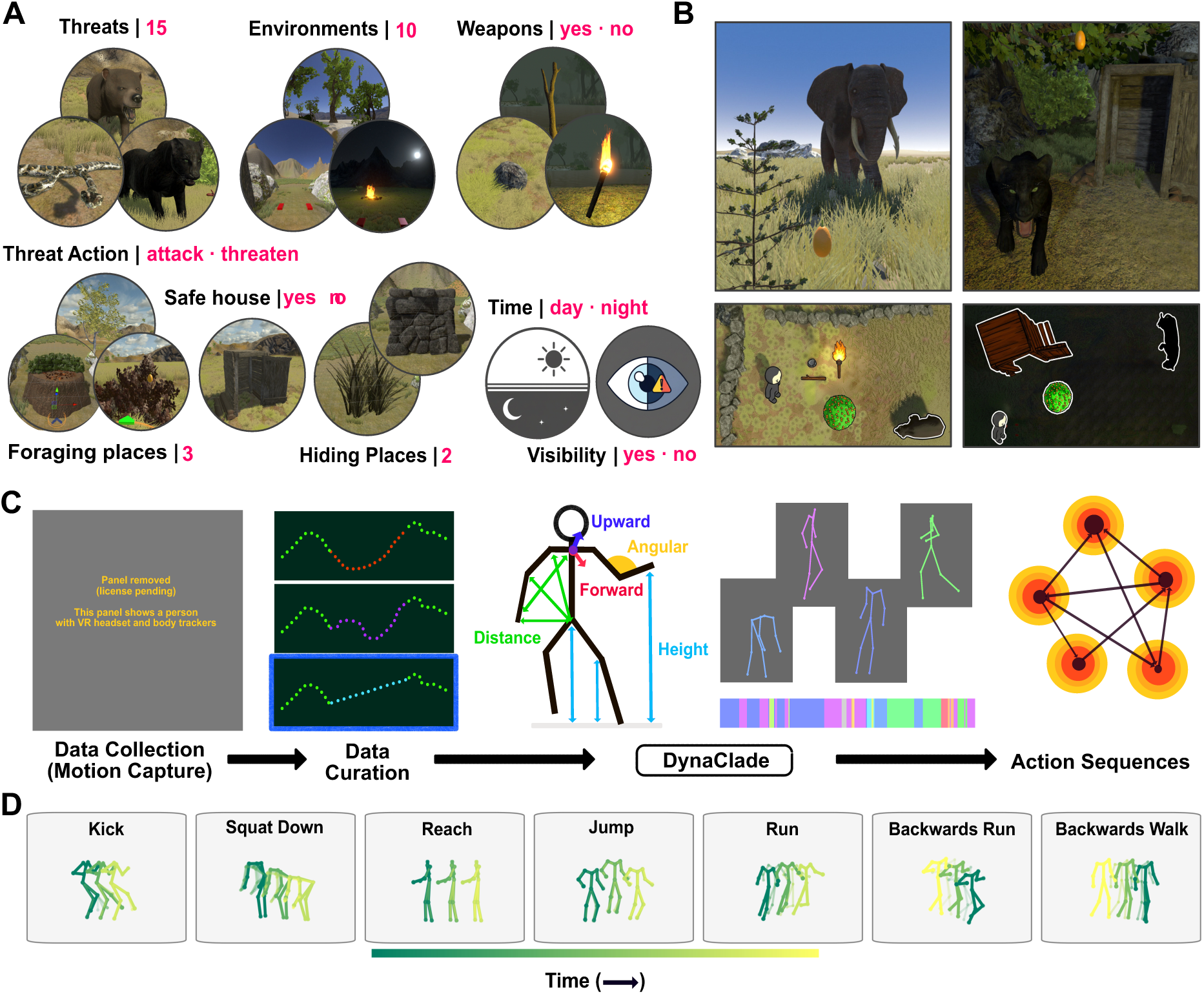
Experimental setup and action segmentation approach. A. Wide range of foraging scenarios designed to permit a rich repertoire of actions. B. Schematic illustration of task elements and configurations. C. DynaClade workflow. D. Illustration of identified actions.

**Figure 2.**
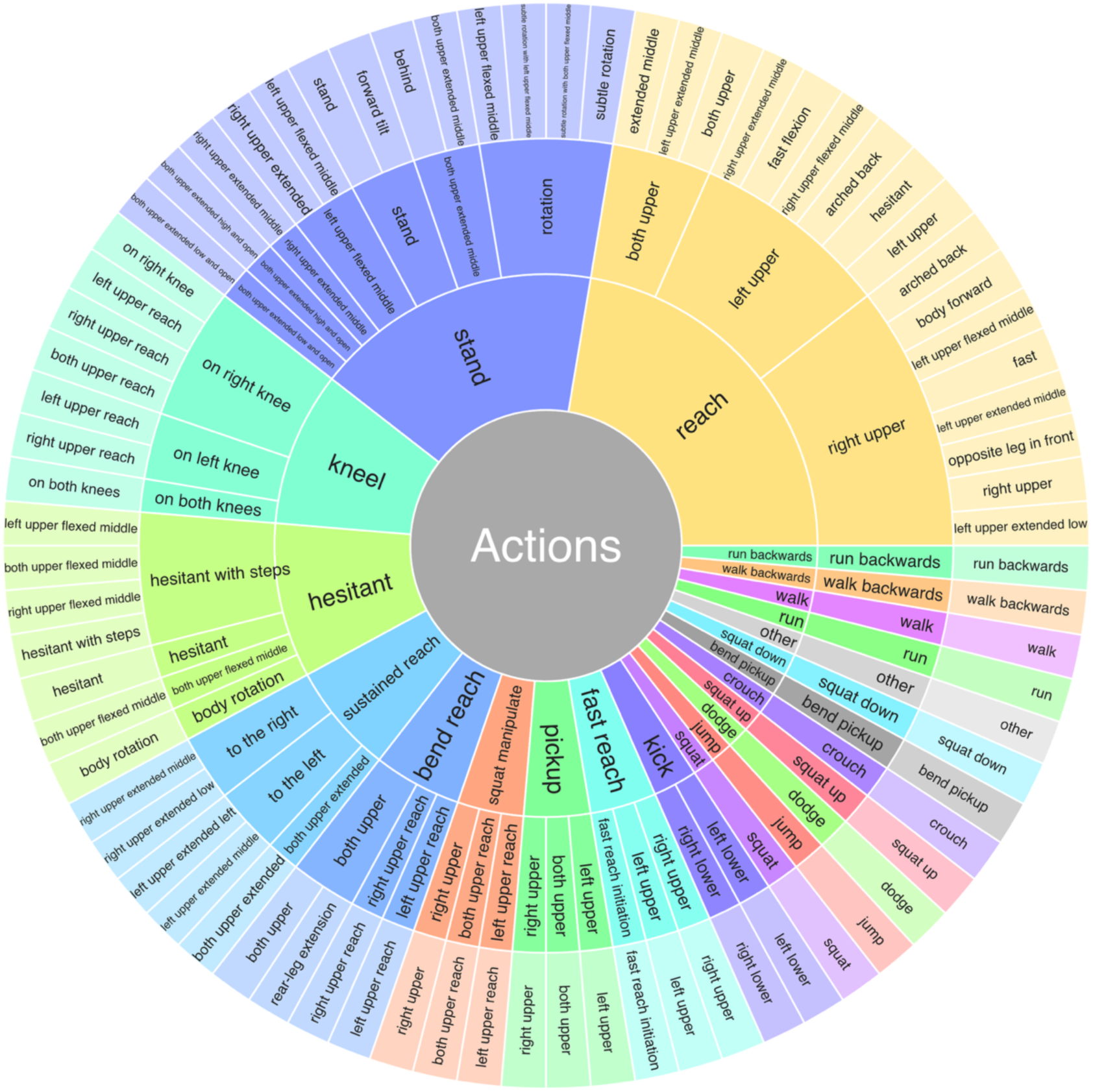
Structure of the identified 22 coarse actions, 46 intermediate actions, and 70 fine actions. Higher-level actions generalise across more specific movement execution styles. An interactive version of this figure is provided in supplemental material.

Because DynaClade is not designed to reproduce human perception of action boundaries, we did not evaluate its accuracy against a human-annotated data set. To evalulate its face validity, we instructed participants in one recording session to perform one of 14 specified action sets (157 epochs in total) and tested whether DynaClade recovered these actions. The predominant action identified for each instruction set corresponded to the instructions, with few exceptions for kinematically difficult or ambiguous actions such as tiptoeing being collapsed with walking (see Supplementary Information for details).

In the following, we report results from the 924 foraging epochs, during which we instructed the goals (collect fruit and avoid contact with threats) but not how to achieve them or which actions to perform. We found that kinematic characteristics of the actions identified by DynaClade were consistent, and their spatial occurrence showed strong correspondence with task affordances. For example, horizontal locomotion speed as assessed from the hips was higher for *run* (2.14 m/s), *jump* (1.04 m/s), and *backward run* (0.94 m/s) than for *backward walk* (0.66 m/s), *forward walk* (0.63 m/s), or for stationary actions such as *stand* (0.12 m/s) or *kneel* (0.03 m/s). Within 1 m of the foraging (fruit collection) position, the most prevalent actions were *reach* (35.4%), *walk* (15.8%), *bend/reach* (9.2%) and *stand* (8.1%), actions that directly reflect the foraging task. Within 1 m of the safe house, the most frequent action was *walk* (63.3%), consistent with escape. Within 1 m of the start position, the most common actions were *stand* (36.3%) and *walk* (32.3%), consistent with instructions at the start of the epoch. Collectively, these results provide evidence for the behavioural validity of the DynaClade approach.

### Characteristics of individual actions

Out of all detected actions, a subset accounted for the vast majority of behaviour. At the three levels of granularity, 11/22 coarse actions, 25/46 intermediate actions, and 37/70 fine actions explained 95% of all time bins. The most frequent actions at the coarse level (figure 3A) were *reach* (29.9%), *walk* (20.0%), *stand* (16.2%), *hesitant* (11.2%), and *bend/reach* (5.1%), and thus primarily focused on foraging and locomotion (figure 3A). Rare but highly specific actions included *dodge*, *backwards run*, *crouch* and *jump* (all < 1%). The average duration of an action bout differed considerably between actions (figure 3A). Foraging-related actions such as *squat/manipulate* (7.7 s), *kneel* (7.6 s), *squat* (3.3 s), and *reach* (2.6 s) lasted longer on average than locomotor (e.g. *walk*, 0.8 s; *run*, 0.5 s) or more specific actions (e.g., *jump*, 0.2 s; *kick*, 0.2 s).

**Figure 3.**
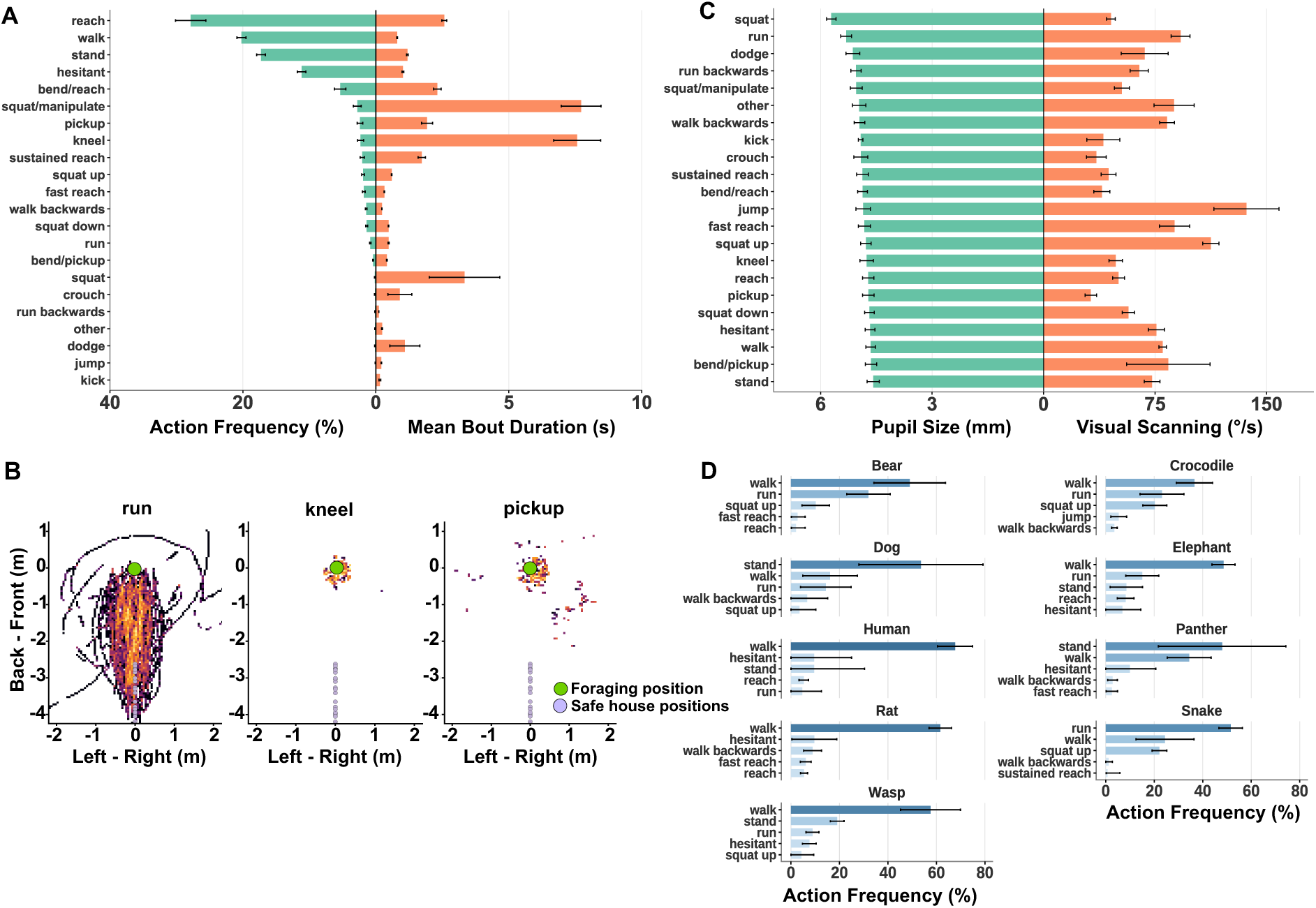
A. Frequency of actions (percent of all time bins) and duration of action bouts. B. Spatial occurence of actions in the different VR scenarios, aligned by foraging position (green dot at coordinate origin) and the position of the safe house (blue dots on negative y-axis), which differed by scenario. C. Pupil size and visual scanning during actions. D. Frequency of escape actions (percent of all escape time bins) for different threats; see methods for detection of escape. Panels ACD show mean ± standard error of participant-level average.

Some actions were highly specific to the task (see examples in figure 3B). For example, *kneel* and *bend/reach* occurred mainly within 0.5 m of the fruit bush (90.3% and 79.1% of overall occurrence, respectively). Fruit collection was executed in four different manners, corresponding to the actions *reach*, *fast reach*, *sustained reach* and *pickup*, with different specificity. While *reach, fast reach* and *pickup* occurred mostly within 0.5 m of the fruit bush (88.9%, 62.1%, and 60.0%, respectively), this percentage was lower for *sustained reach* (44.0%). *Sustained reach* and *pickup* also occurred within 0.5 m of weapon objects that were lying on the ground (30.1% and 25.7%, respectively).

DynaClade identified an action not described in previous work on defensive behaviour, which we labeled *hesitant* after inspection and which comprised 11.2% of all time bins. Hesitant bouts consisted of slow torso movement on the spot, which resembles visual information sampling and is akin to vicarious trial-and-error behaviour in non-human animals (Redish, 2016). Compared to *stand*, *hesitant* was characterised by a higher waist movement speed (0.18 vs. 0.12 m/s, linear mixed-effects model, p < .001, all reported p-values are Holm-Bonferroni corrected for the number of tests), at a longer action bout duration (1.2 s vs. 1.0 s, linear mixed-effects model, p < .001 corrected). Both hesitant and stand were often embedded between walk actions, but this occurred more frequently for hesitant (37%) than for stand (23%, *_χ_*^2^-test, p < .001 corrected). In these cases, both actions entailed a similar change in walking direction (absolute change: hesitant: 91.6°, stand: 94.1°, Watsons two-sample test, n.s.) and a similar bout duration (hesitant: 1.0 s, stand: 0.9 s, linear mixed-effects model, n.s.).

Different actions were associated with different accompanying behaviour, as illustrated by eye tracking data not used by DynaClade for action identification (figure 3C). Pupil size and visual scanning (i.e., cumulative gaze movement) differed widely between actions (linear mixed effects models, both p < .001 corrected), potentially indicating differences in active information sampling and arousal. While pupil size, which is linked to arousal (Reimer et al., 2014), was lowest during *stand*, it was highest during *squat*, *dodge* and *run*. Visual scanning was particularly high during *jump*, *squat up* and *run*, and low during object-oriented actions such as *pickup*, *kick*, and *bend/reach*.

Next, we used this method to ask how people escaped from threat. To identify escape actions, we considered only time periods during which a threat was present in the environment and the participant had not picked up a weapon. We then defined escape as the period between leaving the fruit bush for the last time and entering the safe house. We found that the most prevalent actions during escape were *walk* (47.8%), *run* (21.9%), *squat up* (8.1%), *stand* (5.3%), *hesitant* (4.2%), and *walk backwards* (3.1%). During the 2 s around the interruption of an escape attempt (i.e. when participants initially moved away from the fruit bush and then returned), the most frequent actions were *hesitant* (37.8%) and *walk* (30.6%), followed by *stand* (18.4%). The occurrence of these actions also depended on the threat from which people escaped (figure 3D, linear mixed effects model, p = .017 corrected). For example, when escaping from a snake, the dominant action was *run*. When escaping from a bear or a crocodile, people also ran frequently but less often than when escaping from a snake. Interestingly, when escaping from a panther or a dog, the dominant action was *stand* rather than *walk* or *run* (figure 3D). This may reflect that for these two threats in particular, people often escaped even when not being directly attacked, and stood close by the safe house for some time before entering.

### First-order transitions capture non-random structure

The action vocabulary developed in the previous section describes what participants do, which now allowed us to investigate the organisation of these actions. We began with first-order transitions, that is, the probability of each action given only the one immediately preceding it. As expected, this revealed a highly non-random structure. From 462 possible transitions between two subsequent actions, only 257 (53.1%) occurred in the data, and their distribution was highly non-uniform: the 20 most frequent action bigrams (i.e., action->action transitions) explained more than 70% of transitions.

The most common action bigrams are shown in figure 4A and often included going from stationary to locomotor action. The transition structure is visualised in figure 4B. Information flow analysis of the first-order Markov transition matrix (Rosvall and Bergstrom, 2008) revealed a clear structure with two communities of actions (figure 4B). One community consisted largely of postural elements used for foraging in a stationary position. The other contained foraging actions (*reach*, *fast reach, sustained reach*) as well as locomotion and other dynamic elements. The most common transition between these two communities was from *squat up* (stationary) to *walk* (dynamic). Analysis of the transition matrix further revealed that predictability of the actions increased when participants escaped from a threat: the average conditional entropy of the participant-specific transition structure was reduced from 2.67 bits (all actions) to 2.35 bits (escape sequence, Wilcoxon test, p = .008 corrected).

**Figure 4.**
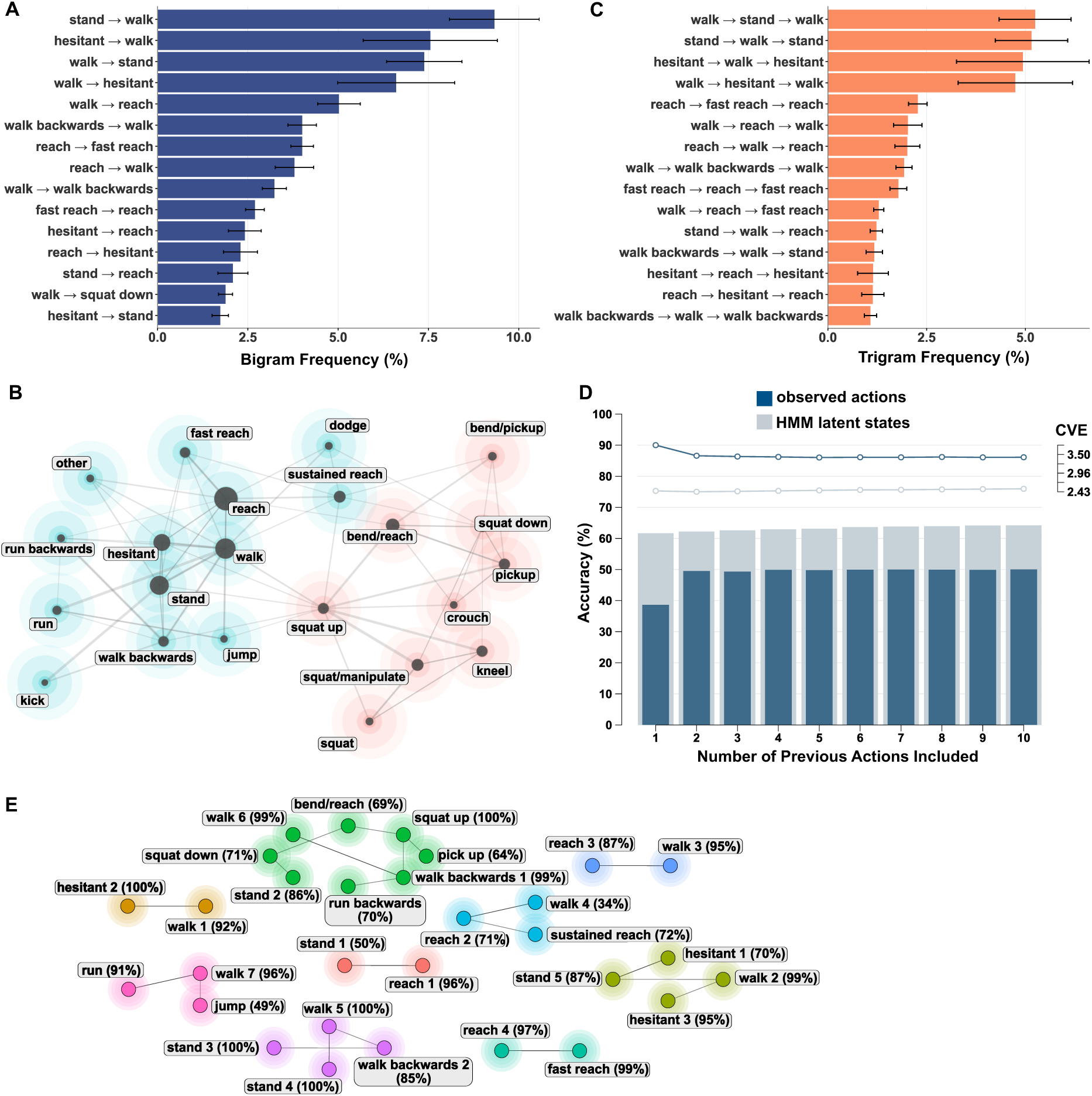
A. Frequency of most frequent action bigrams. B. First-order transition matrix. Node size denotes action frequency, edge thickness transition probability, and the colour bubble denotes two communities of actions identified by information flow analysis. C. Frequency of most frequent action trigrams. D. Ten-fold cross-validated multinomial prediction of the next action, given *n* past actions. Dark bars and line correspond to the observed actions, light bars and line to the latent state in the winning Hidden Markov Model. Left axis shows cross-validated prediction accuracy, right axis shows cross-validated error (CVE). E. Transition structure of latent states in the winning Hidden Markov Model, where edges show the most common transition for each node. Node/bubble colours denote condensation cliques, identified by dominant transitions and thereby indicating action context. Latent states are labelled by their most frequent observed action, and the probability of emitting that action. Error bars in panel A and C: Participant-level standard error of the mean.

### Action sequences have higher-order structure

Next, we sought to disambiguate between the suggestion of stochastic action selection and extended fixed action sequences, by analysing whether action sequences deviated from first-order structure. Importantly, while the potential occurrence of a threat in our task mandated a sequence of abstract goals (foraging fruit, then avoiding or escaping from the threat), the sequence of individual actions to achieve these goals (such as *run*, *walk*, *stand*) was not determined by the task.

Overall, we discovered strong evidence for a non-Markovian sequence structure even within our finite data. Out of 9,702 possible trigrams and 3,468 trigrams expected under the first-order Markov model, only 1,320 occurred in our data, and the 84 most common trigrams explained 70% of the transitions, as opposed to 88 trigrams expected under the first-order model (figure 4C). Notably, none of the most common trigrams (e.g. walk-stand-walk or stand-walk-stand) was directly mandated by the task, underlining that this higher-order structure is not an artefact of our specific task. Next, we tested successively whether *n*-gram frequencies deviated from those predicted by a (*n-1*)-order transition model. We found evidence for an at least 8^th^ order Markov structure in the action sequences (Monte-Carlo permutation test, p = .024 corrected). Notably, this was not simply due to the abstraction inherent in our action hierarchy, since the same result was found for the two finer-grained action levels. When including just the escape sequences, we found evidence for an at least 4^th^ order structure (Monte-Carlo permutation test, p = .014 corrected), based on a smaller number of observations. All of this suggests higher-order structure in the sequences. However, given the large number of observations, this result might be driven by rare sequences. Hence, we were interested in how much variance in the selection of the next action is predicted by previous actions.

To quantify the impact of past actions on the current one, we ran a 10-fold cross-validated multinomial regression, and analysed model error as well as prediction accuracy (figure 4D). Prediction accuracy of current from past actions improved strongly from a first-order (39.6%) to second-order model (49.0%). While model error improved up to a 6^th^-order model, consistent with our in-sample permutation tests, the impact of past actions on model error and prediction accuracy was small beyond the second-order model (improvement in accuracy < 1.0%). This second-order structure was not just due to different participants using different first-order transition models: at the level of single participants, with a substantially smaller amount of data available to fit the models, we observed an increase of 5.0% ± 0.4% (mean ± standard error) in cross-validated prediction accuracy from a first-order model (45.7%) to a second-order model (50.7%).

A common concern in temporal action segmentation is that higher-order structure, and in particular second-order structure, is spuriously induced by mis-classification of single data frames. If, for example, a single frame is randomly flipped, then the next frame can be unambiguously predicted from the one preceding the flipped frame (second-order), but not from the flipped frame itself (first-order model). To account for such mis-classifications suggesting higher-order structure, we fitted a hidden Markov model (HMM) with the same number of states as observations, and analysed the Viterbi-decoded latent state sequences. Again, the decoded sequences exhibited strong evidence for non-Markovian structure of at least 8^th^ order (Monte-Carlo permutation test, p = .020 corrected). Taking all of these results together, we found statistical evidence for a higher-order structure, with most of this structure explained in a second-order model.

### Action context explains higher-order structure

Next, we sought to determine the underlying structure of this higher-order model. We considered different mechanistic hypotheses which we expressed as hidden Markov models (HMM). In our implementation, these HMMs assume a first-order sequence of latent (unobservable) states, which map onto the observed actions in a probabilistic manner. This flexible formalism can account for different latent states leading to the same observed action, but also for the same latent state leading to different observed actions. The first hypothesis we considered was that the combination of an observed action with an abstract goal or task context would define a latent state. For example, walking with the goal of picking fruit and walking with the goal of seeking safety might not be distinguishable in the observed action, but might systematically predict different next actions. If the task or goal context is predicted by the history of preceding observations, then this could lead to a higher-order structure of the observed sequences and thus explain our data. The next hypothesis we tested is that sequences are generated by a hierarchical (or switching) Markov model. Here, observations truly have a first-order structure, but the agent switches between different transition models, for example, from a foraging model to an escape model, as has been suggested in non-human animals (Weinreb et al., 2026). This can be seen as a special case of the first hypothesis and is captured in a HMM where the number of states is a multiple of the number of observations (if all observed states are used in each state of the system). Finally, because of heterogeneity between participants or between scenarios, different first-order models might determine action sequences in different epochs. This could also explain higher-order structure in the data and is captured by mixture HMMs. We fitted all these models to our data. Model selection metrics decisively favoured a HMM with 30 hidden states, compared to 22 observed actions. This supports our first hypothesis, by which observed actions together with action context define a hidden state. While the Viterbi-decoded sequences still demonstrated statistical evidence for an at least 5^th^ order structure of latent states (p = .020 corrected), the first-order model now had 62.2% prediction accuracy, which increased to 63% for the second-order and 65% for higher-order models. This demonstrates that the HMM formalism improves the original model also in terms of explained variance.

The first-order transition structure of the fitted latent states is illustrated in figure 4E, where we label each latent state by its most probable observed state for illustration. Importantly, almost all of the transition probabilities were below 80%, such that no fixed action sequences are expressed in this structure, contrary to one of the theoretical proposals motivating this analysis. To illustrate the community structure of latent states, we sparsified the transition graph to the dominant transitions (i.e., the most probable next state) and considered the ensuing sub-graphs, which we term condensation cliques. While some observed states never constituted a dominant action and others mapped onto one specific latent state, 5 observed actions (*hesitant*, *reach*, *stand*, *walk, walk backwards*) were the dominant action in more than one latent state. Importantly, the multiple instances of these degenerate actions often occurred in separate condensation cliques, or otherwise in different transition contexts. For example, the two latent states associated with *walk backward* occurred in one condensation clique with foraging actions and *run backwards*, and in a second condensation clique containing only *stand* and *walk*. Taken together, these results suggests that latent states in this HMM are characterised by task context together with the emitted action. In other words, within an action context, the precise sequence of actions largely follows a first-order structure, i.e., there is no evidence for a substantial prevalence of fixed action sequences beyond the first order.

## Discussion

In this work, we use motion capture data from an ecologically realistic fm-VR task, and propose the unsupervised action segmentation algorithm DynaClade, to understand how humans organise sustained streams of actions in a risky foraging task. We find that DynaClade discovers a meaningful action repertoire including a hitherto undescribed action which we label *hesitant* and which is characterised by slow torso movements on the spot. The action repertoire is validated by correspondence to specific instructions in instructed epochs as well as with environmental features in foraging epochs, by kinematic plausibility, and by systematic differences in eye-tracking and pupil size metrics. We then analyse action sequences and find that they are largely organised in a first-order Markov chain of context-dependent latent states.

Understanding how humans organise sustained streams of spontaneous actions to achieve their goals is a central challenge in behavioural science, as this constitutes the majority of daily life activities (Diedrichsen et al., 2026). Various theoretical frameworks have been proposed for the assembly of action sequences. Reflective planning appears to be the optimal solution to many action selection problems (Mattar and Lengyel, 2022), but this becomes quickly intractable even in relatively small state spaces (Daw et al., 2005). Non-exhaustive approximations that work well in structured domains, such as Monte Carlo tree search, can saturate a system’s computational capacity in even slightly extended temporal horizons (Espinosa et al., 2022; Brochard et al., 2025). Some theoretical proposals have focused on reducing planning granularity, that is, prospectively planning abstract goals and leaving motor execution details to subordinate controllers in a hierarchical system (Merel et al., 2019). An alternative is to reduce the depth of the planning time horizon (Brochard et al., 2025; Diedrichsen et al., 2026), such that actions are selected only shortly before they are executed, shaped by the history of past actions. This could be implemented by reflective planning as well as by reflexive, e.g. habitual, action selection. Even more simply, the next action could be selected by considering only the current action, according to a stochastic model that considers goals and context. This is what has been proposed for human action selection based on analyses of behavioural lapses (Cooper and Shallice, 2000, 2006; Schwartz, 2006). Our data support this view with an entirely different method, by analysing spontaneous naturalistic behaviour in a self-guided task with limited structure. Thus, humans appear to use a computationally cheap and efficient strategy to render the assembly of self-guided behaviour tractable. While spontaneous fixed action sequences have been informally observed in humans, this could be due to requirements of external tasks, particularly in the interaction with technical equipment. Thus, it remains an interesting question how humans deal with the increased computational complexity of prospective planning in more structured environments than the one presented here.

Our observation that human action selection is largely guided just by the last action is consistent with an observation from rodent behaviour (Wiltschko et al., 2015; Weinreb et al., 2026). However, the MoSeq algorithm underlying these non-human results assumed a first-order Markov structure a priori, different from the DynaClade algorithm. Thus, there is a possibility that mouse action sequences, or the residual sequences after model fitting, would exhibit higher-order structure, too. On the level of the fitted models, the main difference between rodent and our human results is that mice appear to switch between different first-order models that exhaust the entire action repertoire (Weinreb et al., 2026), whereas in our data, different action contexts utilised only a small part of the action repertoire. Whether this is due to the smaller amount of data available to us, the different structure of the environment – with mice behaving spontaneously and humans following instructed high-level goals –, or a genuine difference between species, remains a question for future work.

Our action repertoire captured not only locomotor and foraging behaviours but also actions whose functional significance was not explicitly prescribed by the task. Among these, an interesting action is what we labelled as *hesitant*. Hesitant was characterised by slow body sway on the spot, and occurred frequently between locomotor bouts in a manner suggestive of decision points. Hesitant behaviour has been described in non-humans at decision points (Redish, 2016; Geramita et al., 2026), but has not been kinematically characterised in humans. Importantly, it seems to be distinct from freezing behaviour, which has occasionally been reported in humans and is characterised by decreased, rather than increased, body sway (Roelofs and Dayan, 2022).

An important open question concerns the state space of action selection from the perspective of the agent. We have formulated our results in terms of discrete action units, in line with a large and growing literature (Ding et al., 2024). However, it remains unclear whether this is only a useful abstraction from an observer’s perspective, or actually constitutes the substrate of action selection in the nervous system, as some non-human work suggests (Markowitz et al., 2018, 2023). Alternative action selection frameworks have been proposed at the level of continuous action affordances (Pezzulo and Cisek, 2016). In a dynamical-systems view of the brain (Durstewitz et al., 2023; Churchland and Shenoy, 2024), the two ideas are not mutually exclusive, since dynamical systems contain discrete elements such as modes and attractors. However, at present we are lacking a coherent and empirically grounded framework to map one perspective onto the other. HMMs have in fact been suggested as means to integrate neural and behavioural data into discrete states (Dommanget-Kott et al., 2026).

DynaClade successfully extracted a valid, hierarchically structured action repertoire from complex, naturalistic full-body motion capture data, without presupposing action identity or sequence structure. Alternative action segmentation approaches are typically based on the assumption of a first-order sequence structure (Wiltschko et al., 2015; Weinreb et al., 2024), or an a-priori definition of the action space (Ding et al., 2024). DynaClade shares the general approach of B-SOID (Hsu and Yttri, 2021) and A-SOID (Tillmann et al., 2024) and renders it suitable for our data domain. Nonetheless, several extensions would broaden the applicability of DynaClade, for example a capability for handling incomplete data (Rose et al., 2026), and for operating on generic body models that can be applied to other data sets without retraining. Importantly, there is compelling evidence that instructed action sequences are represented and generated in a temporally hierarchical manner (Yokoi and Diedrichsen, 2019; Eckstein and Collins, 2021), such that future action segmentation work might focus on extracting this temporal hierarchy.

To summarise, we demonstrate that human behaviour in ecologically valid settings is organised in a rich repertoire of actions whose sequential assembly is largely explained by context-specific first-order Markov processes. These findings reveal a level of sequential organisation in human behaviour that is inaccessible to conventional laboratory paradigms, and establish computational ethology as a productive framework for characterising the structure, function, and individual variability of naturalistic human action.

## Methods

### Participants

We recruited 30 participants (17 female, 13 male; mean age ± SD = 22.4 ± 3.22 years) from the student and general population. The study was conducted in accordance with the Declaration of Helsinki and approved by the Research Ethics Committee of the Medical Faculty, University of Bonn (240/22). All participants provided written informed consent before the experiment and received a fixed monetary compensation for their participation. As in previous work using a similar setup (Sporrer et al., 2023; Hutabarat et al., 2025; Kornemann et al., 2026b, 2026a), potential participants were pre-screened and excluded if they reported a history of life-threatening interpersonal trauma, any life-threatening trauma and current symptoms of post-traumatic stress disorder as assessed using the PCL-5 checklist (Weathers et al., 2013), or any diagnosed mental health condition (including substance abuse) or neurological disorder. To minimise risk of accidents, individuals with self-reported impairments to movement, vision, or hearing were also excluded. Kinematic summary statistics and self-report data from this sample were included in a previous psychological investigation (Kornemann et al., 2026a).

### Experimental procedure

We sought to create a comprehensive data set of behaviour in biologically relevant virtual environments. To this end, participants performed a foraging task (collecting fruit on various trees/bushes) in fm-VR, during which they could encounter a threatening object, animal or human. Participants were explicitly instructed to collect as many fruit as possible, while avoiding physical contact with any of the threats. There were no further instructions, including no instructions how to achieve these goals. The experiment was structured into foraging epochs that typically lasted around 30 s but could be considerably shorter or longer (up to 90 s), depending on the participant’s behaviour. During each epoch, a threat could appear, and participants were free to react naturally. To give participants experience with the VR settings, the first epoch lasted 180 s and participants were explicitly instructed that no threat would appear in this epoch, and that they should explore the environment.

Each foraging epoch implemented one of 80 unique scenarios that were systematically designed to elicit a broad range of movements and maintain engagement (Table S1). This was achieved by varying environments (12 distinct natural scenes), interactive objects such as different fruit collection plants (bush or tree with fruit appearing in low, medium or high position), barriers, hiding spots, safe house, weapons (sticks, stones, torches), and threats (human, panther [black-coated leopard], crocodile, brown bear, African elephant, dog [Dobermann], snake [viper], rat, spider, wasp swarm, scorpion, dynamite, falling rock). The diversity of human, non-human animal, and inanimate threats was based on previous work suggesting that threat size, attack style and threat category strongly influence participants’ escape behaviour (Sporrer et al., 2023). The safe house was automatically closed when fully entered as long as the threat was not already blocking the door. Weapons, when used against a threat, had a 1/3 chance of killing the threat, stopping its attack, or being ineffective. All elements were combined into biologically plausible combinations, i.e. we sought to place a certain threat only into a natural scene in which it might reasonably be encountered. An epoch would end by closing the safe house, killing the threat, getting killed by the threat, or by surviving for a pre-defined maximum duration. To demonstrate the realism and immersiveness of the VR epochs, we analysed participants’ spontaneous vocalisations in line with our previous work (Sporrer et al., 2023) and found threat-directed vocalisations on 7.6% of trials and in 60.0% of participants, including screams on 0.4% of trials and in 13.3% of participants. Participants also followed instructions, collecting (mean ± SD) 12.5 ± 11.5 fruit, and virtually surviving on 77.3% of trials. Because of equipment malfunction, not all participants completed all epochs. Overall, 2341 foraging epochs were completed per protocol.

After this main experiment, participants were instructed in a second session to perform one out of 14 specific actions (e.g., walk, crawl, jump, punch; Table S1). This served to provide ground-truth data for the validation of our action segmentation algorithm. Overall, 364 instructed epochs were completed per protocol.

### Software and equipment

The experiment was created and run using the VRthreat toolkit for Unity (Brookes et al., 2023) and the Unity Experiment Framework (UXF, Brookes et al., 2020) in the Unity game engine (Unity Technologies, San Francisco, CA, USA) together with Steam VR (Valve Corporation, Bellevue, WA, USA). We used a wireless HTC Vive Pro Eye headset (HTC Corporations, Taoyuan, Taiwan; 1440×1600 pixels per eye, 90 Hz). Data collection took place in a 7.5 m x 17 m physical space at the University of Bonn.

Full body participant movements were recorded using 10 Qualisys Arqus A9 cameras at 200 Hz with Qualisys Track Manager (v2023.3). Marker placement followed the 41-point sports marker set by Qualisys Track Manager, a derivative of the conventional gait model, to define anatomical landmarks and segment axes. Participant-specific Automatic Identification Models (AIM) were generated and calibrated using both a static A-pose trial and a short dynamic trial to account for individual anthropometric and marker placement variations.

#### Motion capture data pre-processing

Raw marker trajectories were processed within Qualisys Track Manager. Automated labelling and gap-filling was performed using linear and rigid body interpolation. The pre-processed trajectories were then passed through an inverse kinematics solver to reconstruct the skeletal pose. This procedure calculated the 3D joint centre location represented in 3D global positions and orientations represented as unit quaternions.

#### Quality Control

To ensure data integrity, a multi-step quality check was performed on an epoch-by-epoch basis. An epoch was included for further analysis if it passed three criteria. First, data completeness was verified, requiring at least 95% of data points to be present for each tracked joint. Second, anatomical consistency was checked by ensuring all joint-to-joint distances remained within a 10% deviation for at least 95% of the epoch’s duration. Finally, a visual inspection was conducted to confirm that the recorded movements were anatomically feasible. After quality control, 1084 epochs from 23 participants were retained for DynaClade. In three foraging epochs, the VR task had been interrupted (i.e. not delivered per protocol), so that participants’ behaviour did not correspond to the VR environment; these epochs were used for action identification with DynaClade but not analysed further. For one foraging epoch, Unity output was corrupted due to a technical malfunction; this epoch was used for action identification and analysis of action frequencies, but is excluded from analyses relating to spatiotemporal task features. Analysis of the identified actions is thus based on 1081 (924 foraging and 157 instructed epochs).

#### Data Smoothing

The validated data were smoothed to attenuate high-frequency noise resulting from the capture process. This was achieved using a rolling median filter with a window size of 160 ms (16 data points at 100 Hz sampling rate) followed by a Butterworth low-pass filter with a cutoff frequency of 6 Hz. The parameters for both filters were fine-tuned to remove noise without distorting the underlying movement signal.

### Temporal action segmentation

Temporal action segmentation aims to divide continuous movement into distinct behavioral units and identify their temporal boundaries. However, what constitutes a behavioral unit differs across scientific fields and levels of behavioral abstraction. In computer vision, temporal action segmentation typically focuses on semantically defined actions within longer activities, whereas computational ethology often seeks to identify shorter, recurring movement patterns, commonly termed behavioral motifs or syllables (Berman, 2018; Ding et al., 2024). Regardless of their granularity, these behavioral units may either be predefined or discovered from the data, corresponding to supervised and unsupervised learning approaches, respectively. Supervised methods learn to identify predefined actions from annotated examples, while unsupervised methods seek to discover recurring behavioral units without prior action labels. The latter are particularly relevant when the behavioral repertoire is not known in advance, as in the present work.

A common approach to unsupervised behavioral discovery involves representing high-dimensional movement data in a lower-dimensional space and identifying recurring patterns through clustering. Following this framework, B-SOiD (Hsu and Yttri, 2021), developed for animal behavior analysis, extracts spatiotemporal features from tracked body parts, projects them into a lower-dimensional space using Uniform Manifold Approximation and Projection (UMAP) and identifies behavioral clusters using HDBSCAN. Alternatively, behavioral states and their temporal dynamics can be modeled within a probabilistic framework. MoSeq (Wiltschko et al., 2015) and Keypoint-MoSeq (Weinreb et al., 2024), employ an autoregressive hidden Markov model (AR-HMM), in which each latent behavioral state is characterized by distinct movement dynamics, while transitions between states follow a first-order Markov process.

While MoSeq has been successfully used to investigate mouse behaviour (Markowitz et al., 2018, 2023; Weinreb et al., 2026), its explicit assumptions about movement dynamics and state transitions may not generalize to more complex human actions. We therefore chose to build on B-SOiD, adapting its clustering-based approach to 3D human motion capture data, which differed significantly from the regime of 2D (mainly non-human) motion capture in which B-SOID was validated. However, applying B-SOiD to our recordings resulted in very short behavioral motifs, fragmenting longer, coherent actions that were readily identifiable by human observers. We reasoned that this fragmentation was primarily due to B-SOiD’s choice of dynamic features, which were optimized for mouse movements and might not adequately capture the dynamics of human actions. Thus, we complemented the algorithm with a much larger feature space that included higher-order derivatives, thus providing a more succinct description of the action dynamics on a frame-by-frame level. Furthermore, we added a human-in-the-loop postprocessing step to ensure the functional coherence of the identified action classes. Thus, we developed the DynaClade algorithm which comprises 5 steps.

#### Step 1: Feature space

To provide a rich, quantitative description of human movement, we extracted a high-dimensional kinematic feature set from the raw 3D motion capture data. This process was designed to capture both the instantaneous postural configuration of the body and its dynamic evolution over time on a frame-by-frame level. The final feature set comprised 311 distinct kinematic variables for each time point, calculated as follows.

A set of **77 base features** was first computed to describe the participant’s spatial posture and orientation:

- Postural Configuration Features (75 features):

- **Inter-Joint Distances (55 features):** We calculated the Euclidean distances between all 55 pairs of non-adjacent anatomical markers. These features encode the overall shape and configuration of the body, capturing relationships between limbs and the torso.
- **Joint Heights (12 features):** The vertical distance (z-axis) from 12 key joints to the floor plane was computed. These features are critical for identifying changes in elevation, such as crouching, jumping, or bending.
- **Segmental Angles (8 features):** We calculated 8 key intra-limb angles, such as the flexion/extension of the elbows and knees, to describe the articulation of individual body segments.
- Global Body Orientation Features (2 features):

- **Torso Inclination (1 feature):** The angle of the torso vector relative to the horizontal floor plane was calculated to quantify forward or backward bending.
- **Torso Orientation (1 feature):** The yaw of the torso was calculated as the angle of its forward-facing vector projected onto the horizontal plane, indicating the participant’s heading direction.

To capture the motion dynamics, we then computed the **temporal derivatives** for each of the 77 base features. This yielded the instantaneous **velocity** (first derivative), **acceleration** (second derivative), and **jerk** (third derivative) for each spatial feature, adding **231 dynamic features** (77 base features × 3 derivatives) to our set.

Finally, to represent the overall translational movement of the participant’s centre of mass, we calculated three **global kinematic features**:

- Global Velocity, Acceleration, and Jerk (3 features) of the participant’s hip centre marker.

#### Step 2: Dimensionality Reduction with UMAP

Next, we projected these features into a lower-dimensional embedding that preserves the data’s topological structure. Prior to dimensionality reduction, all features were z-transformed. We then employed Uniform Manifold Approximation and Projection (UMAP version 0.5.7; McInnes et al., 2020; Healy and McInnes, 2024), to create the data embedding. The target number of dimensions for this embedding was determined by a preliminary Principal Component Analysis (PCA), in which 84 components were required to explain 95% of the data variance. The UMAP algorithm was configured with n_neighbors=300, min_dist=0.0, and densemap=False.

#### Step 3: Unsupervised Clustering with HDBSCAN

Following dimensionality reduction, we identified initial clusters within the UMAP embedding using Hierarchical Density-Based Spatial Clustering of Applications with Noise (HDBSCAN, Campello et al., 2013, 2015), (version 0.8.39). Hyperparameters for the model were tuned using a grid search, and relative validation was performed using a metric based on internal cluster cohesion and separation [33, 34, 35]. The final HDBSCAN model was configured with the following parameters: min_cluster_size=300, min_samples=None, allow_single_cluster=True, and approx_min_span_tree=False.

#### Step 4: Human-in-the-loop postprocessing of cluster centroids

The initial clustering process identified several hundred distinct clusters. To render these data-driven clusters interpretable and to consolidate functionally similar movements, we performed a manual labelling and merging process. A representative prototype for each cluster was generated by selecting data points with the highest membership probability scores from the HDBSCAN output. These prototypes were visually analysed, and clusters representing visually similar movements were merged. Finally, to create a structured and multi-level understanding of the observed behaviours, a three-level hierarchical dictionary was constructed by grouping the refined actions into broader parent categories (e.g., “right kick” and “left kick” actions were grouped into a parent “kick” category). This hierarchy allows for analysis at multiple spatial granularities.

#### Step 5: Classification of ambiguous actions

The initial discovery pipeline leaves some time points unclassified (as noise). To create an efficient method for assigning a final action label to every time point, we trained a supervised classification model. A Random Forest classifier (n_estimators=100) was implemented in Python using the scikit-learn library (Pedregosa et al., 2011). Crucially, this model was trained using the original high-dimensional feature set as input, with the target labels being the final, human-curated action labels derived from the merging and labelling process. The trained classifier was then used to predict the most likely actions for all data points originally classified as noise by HDBSCAN, resulting in contiguous action sequences for the final analysis.

### Statistical analysis

Action sequences and movement trajectories were analysed in R (version 4.5.3) via RStudio (version 2026.08.1 build 195). All code including a list of the required packages and their versions will be available on OSF. Linear mixed effects models were computed using the package lmerTest (Kuznetsova et al., 2017). Tests of Markov structure were computed using successive Monte-Carlo permutation tests as implemented in the package Nestimate (Saqr et al., 2026). To predict future actions from past ones, we implemented a 10-fold cross-validated multinomial model in the package glmnet. Hidden Markov Models were fit with the package seqHMM (Helske and Helske, 2019) and selected based on Akaike Information Criterion. To account for the multiple comparison problem, all statistical tests were Holm-Bonferroni corrected for the number of tests performed, and we report corrected *p*-values in the results.

## Code and data availability

The VRthreat toolkit for Unity is available under an academic license from XIP (https://xip.uclventures.com/product/vrthreat-toolkit-for-unity). The Unity Build used in this study is available as an executable on OSF (https://osf.io/uzghp). DynaClade is available on GitHub (https://github.com/userratos/DynaClade). All R code will be made available on OSF. Curated data will be made available on OSF. Source data are available from the corresponding author under a data protection agreement in line with local ethics regulations.

## References

Anderson DJ, Perona P (2014) Toward a Science of Computational Ethology. Neuron 84:18–31.

Bach DR (2022) Cross-species anxiety tests in psychiatry: pitfalls and promises. Mol Psychiatry 27:154– 163.

Bach DR, Moutoussis M, Bowler A, Neuroscience in Psychiatry Network consortium, Dolan RJ (2020) Predictors of risky foraging behaviour in healthy young people. Nat Hum Behav 4:832–843.

Berman GJ (2018) Measuring behavior across scales. BMC Biol 16:23.

Botvinick MM (2008) Hierarchical models of behavior and prefrontal function. Trends in Cognitive Sciences 12:201–208.

Brochard J, Dayan P, Bach DR (2025) Critical intelligence: Computing defensive behaviour. Neuroscience & Biobehavioral Reviews 174:106213.

Brookes J, Hall S, Frühholz S, Bach DR (2023) Immersive VR for investigating threat avoidance: The VRthreat toolkit for Unity. Behav Res Available at: https://link.springer.com/10.3758/s13428-023-02241-y [Accessed December 12, 2023].

Brookes J, Warburton M, Alghadier M, Mon-Williams M, Mushtaq F (2020) Studying human behavior with virtual reality: The Unity Experiment Framework. Behav Res 52:455–463.

Campello RJGB, Moulavi D, Zimek A, Sander J (2013) A framework for semi-supervised and unsupervised optimal extraction of clusters from hierarchies. Data Min Knowl Disc 27:344– 371.

Campello RJGB, Moulavi D, Zimek A, Sander J (2015) Hierarchical Density Estimates for Data Clustering, Visualization, and Outlier Detection. ACM Trans Knowl Discov Data 10:1–51.

Churchland MM, Shenoy KV (2024) Preparatory activity and the expansive null-space. Nat Rev Neurosci 25:213–236.

Cooper R, Shallice T (2000) CONTENTION SCHEDULING AND THE CONTROL OF ROUTINE ACTIVITIES. Cognitive Neuropsychology 17:297–338.

Cooper RP, Shallice T (2006) Hierarchical schemas and goals in the control of sequential behavior. Psychological Review 113:887–916.

Daw ND, Niv Y, Dayan P (2005) Uncertainty-based competition between prefrontal and dorsolateral striatal systems for behavioral control. Nat Neurosci 8:1704–1711.

Diedrichsen J, Kashefi M, Nazerzadeh A, Pruszynski JA (2026) Online planning of sequential actions. Trends in Cognitive Sciences:S1364661326001257.

Ding G, Sener F, Yao A (2024) Temporal Action Segmentation: An Analysis of Modern Techniques. IEEE Trans Pattern Anal Mach Intell 46:1011–1030.

Dommanget-Kott M, Fernandez-de-Cossio-Diaz J, Coraggioso M, Bormuth V, Monasson R, Debrégeas G, Cocco S (2026) Linking brain and behavior states in Zebrafish Larvae locomotion using hidden Markov models Louis M, ed. PLoS Comput Biol 22:e1013762.

Durstewitz D, Koppe G, Thurm MI (2023) Reconstructing computational system dynamics from neural data with recurrent neural networks. Nat Rev Neurosci 24:693–710.

Eckstein MK, Collins AGE (2021) How the Mind Creates Structure: Hierarchical Learning of Action Sequences. Cogsci 43:618–624.

Espinosa G, Wink GE, Lai AT, Dombeck DA, MacIver MA (2022) Achieving mouse-level strategic evasion performance using real-time computational planning. Available at: http://arxiv.org/abs/2211.02700 [Accessed March 14, 2025].

Geramita MA, Ahmari SE, Yttri EA (2026) The striatal indirect pathway mediates hesitation. Nat Neurosci 29:287–292.

Gökay U, Spurio F, Bach DR, Gall J (2025) Skeleton Motion Words for Unsupervised Skeleton-Based Temporal Action Segmentation. Proceedings of the IEEE/CVF International Conference on Computer Vision (ICCV):12101–12111.

Healy J, McInnes L (2024) Uniform manifold approximation and projection. Nat Rev Methods Primers 4:82.

Helmich RC, Aarts E, De Lange FP, Bloem BR, Toni I (2009) Increased Dependence of Action Selection on Recent Motor History in Parkinson’s Disease. J Neurosci 29:6105–6113.

Helske S, Helske J (2019) Mixture Hidden Markov Models for Sequence Data: The seqHMM Package in R. Journal of Statistical Software 88:1–32.

Hsu AI, Yttri EA (2021) B-SOiD, an open-source unsupervised algorithm for identification and fast prediction of behaviors. Nat Commun 12:5188.

Hunt LT, Daw ND, Kaanders P, MacIver MA, Mugan U, Procyk E, Redish AD, Russo E, Scholl J, Stachenfeld K, Wilson CRE, Kolling N (2021) Formalizing planning and information search in naturalistic decision-making. Nat Neurosci 24:1051–1064.

Hutabarat Y, Sporrer JK, Brookes J, Zabbah S, Kornemann L, Domenici P, Bach DR (2025) Human escape follows a structured movement pattern shaped by threat and context. Available at: http://biorxiv.org/lookup/doi/10.1101/2025.07.27.662958 [Accessed August 10, 2025].

Janssen M, LeWarne C, Burk D, Averbeck BB (2022) Hierarchical Reinforcement Learning, Sequential Behavior, and the Dorsal Frontostriatal System. J Cogn Neurosci 34:1307–1325.

Kornemann L, de Vries O, Hutabarat Y, Serratos Hernandez UD, Bach DR (2026a) Feelings and defensive actions are dissociated under naturalistic threat. Available at: osf.io/preprints/psyarxiv/kxpjm_v1.

Kornemann L, Hutabarat Y, Liu H, Pensel MC, Serratos Hernandez UD, Sporrer JK, Zabbah S, Bach DR (2026b) Running in Tight Spaces: A Large-scale Assessment of Physical and Psychological Safety Risks in Freely Moving Virtual Reality. Available at: osf.io/preprints/psyarxiv/7cqvx_v1.

Krakauer JW, Ghazanfar AA, Gomez-Marin A, MacIver MA, Poeppel D (2017) Neuroscience Needs Behavior: Correcting a Reductionist Bias. Neuron 93:480–490.

Kuznetsova A, Brockhoff PB, Christensen RHB (2017) lmerTest Package: Tests in Linear Mixed Effects Models. Journal of Statistical Software 82:1–26.

Markowitz JE, Gillis WF, Beron CC, Neufeld SQ, Robertson K, Bhagat ND, Peterson RE, Peterson E, Hyun M, Linderman SW, Sabatini BL, Datta SR (2018) The Striatum Organizes 3D Behavior via Moment-to-Moment Action Selection. Cell 174:44–58 e17.

Markowitz JE, Gillis WF, Jay M, Wood J, Harris RW, Cieszkowski R, Scott R, Brann D, Koveal D, Kula T, Weinreb C, Osman MAM, Pinto SR, Uchida N, Linderman SW, Sabatini BL, Datta SR (2023) Spontaneous behaviour is structured by reinforcement without explicit reward. Nature 614:108–117.

Mattar MG, Lengyel M (2022) Planning in the brain. Neuron 110:914–934.

McInnes L, Healy J, Melville J (2020) UMAP: Uniform Manifold Approximation and Projection for Dimension Reduction. Available at: http://arxiv.org/abs/1802.03426 [Accessed April 29, 2026].

Merel J, Botvinick M, Wayne G (2019) Hierarchical motor control in mammals and machines. Nat Commun 10:5489.

Mobbs D, Kim JJ (2015) Neuroethological studies of fear, anxiety, and risky decision-making in rodents and humans. Curr Opin Behav Sci 5:8–15.

Mobbs D, Wise T, Suthana N, Guzmán N, Kriegeskorte N, Leibo JZ (2021) Promises and challenges of human computational ethology. Neuron 109:2224–2238.

Pedregosa F, Varoquaux G, Gramfort A, Michel V, Thirion B, Grisel O, Blondel M, Prettenhofer P, Weiss R, Dubourg V, Vanderplas J, Passos A, Cournapeau D, Brucher M, Perrot M, Duchesnay É (2011) Scikit-learn: Machine Learning in Python. Journal of Machine Learning Research 12:2825– 2830.

Pezzulo G, Cisek P (2016) Navigating the Affordance Landscape: Feedback Control as a Process Model of Behavior and Cognition. Trends Cogn Sci 20:414–424.

Reddy G, Desban L, Tanaka H, Roussel J, Mirat O, Wyart C (2022) A lexical approach for identifying behavioural action sequences Faisal AA, ed. PLoS Comput Biol 18:e1009672.

Redish AD (2016) Vicarious trial and error. Nat Rev Neurosci 17:147–159.

Reimer J, Froudarakis E, Cadwell CR, Yatsenko D, Denfield GH, Tolias AS (2014) Pupil fluctuations track fast switching of cortical states during quiet wakefulness. Neuron 84:355–362.

Roelofs K, Dayan P (2022) Freezing revisited: coordinated autonomic and central optimization of threat coping. Nat Rev Neurosci 23:568–580.

Rose F, Michaluk M, Blindauer T, Ignatowska-Jankowska BM, O’Shaughnessy L, Stephens GJ, Pereira TD, Uusisaari MY, Bozek K (2026) Deep Imputation for Skeleton data (DISK) for behavioral science. Nat Methods 23:236–247.

Rosvall M, Bergstrom CT (2008) Maps of random walks on complex networks reveal community structure. Proc Natl Acad Sci USA 105:1118–1123.

Saqr M, López-Pernas S, Misiejuk K (2026) Nestimate: An R Package for dynamic, probabilistic, and high order network analysis. Appl Psychol Meas:01466216261472019.

Schoener TW (1971) Theory of feeding strategies. Annual review of ecology and systematics:369–404.

Schwartz MF (2006) The cognitive neuropsychology of everyday action and planning. Cognitive Neuropsychology 23:202–221.

Sporrer JK, Brookes J, Hall S, Zabbah S, Serratos Hernandez UD, Bach DR (2023) Functional sophistication in human escape. iScience 26:108240.

Tillmann JF, Hsu AI, Schwarz MK, Yttri EA (2024) A-SOiD, an active-learning platform for expert-guided, data-efficient discovery of behavior. Nat Methods 21:703–711.

Trier HA, O’Reilly JX, Spiering L, Yishan SM, Kolling N, Rushworth MFS, Scholl J (2025) Emotions and individual differences shape human foraging under threat. Nat Mental Health 3:444–465.

Verwey WB (2024) C-SMB 2.0: Integrating over 25 years of motor sequencing research with the Discrete Sequence Production task. Psychon Bull Rev 31:931–978.

Weathers FW, Litz BT, Keane TM, Palmieri PA, Marx BP, Schnurr PP (2013) The PTSD Checklist for DSM-5 (PCL-5) – LEC-5 and Extended Criterion A. Available at: https://www.ptsd.va.gov/professional/assess-ment/adultsr/ptsd-checklist.asp.

Weinreb C, Kannan LT, Newman-Boulle A, Sainburg T, Gillis WF, Plotnikoff A, Makowska S, Pearl JE, Osman MAM, Linderman SW, Datta SR (2026) Spontaneous behavior is a succession of self-directed tasks. Neuron 114:922–937.e12.

Weinreb C, Pearl JE, Lin S, Osman MAM, Zhang L, Annapragada S, Conlin E, Hoffmann R, Makowska S, Gillis WF, Jay M, Ye S, Mathis A, Mathis MW, Pereira T, Linderman SW, Datta SR (2024) Keypoint-MoSeq: parsing behavior by linking point tracking to pose dynamics. Nat Methods 21:1329–1339.

Wiltschko AB, Johnson MJ, Iurilli G, Peterson RE, Katon JM, Pashkovski SL, Abraira VE, Adams RP, Datta SR (2015) Mapping Sub-Second Structure in Mouse Behavior. Neuron 88:1121–1135.

Yokoi A, Diedrichsen J (2019) Neural Organization of Hierarchical Motor Sequence Representations in the Human Neocortex. Neuron 103:1178–1190.e7.

