## Supplementary information for "The Computational Grammar of Human Behaviour in a Biological Environment"

### Supplemental material

### Analysis of instructed actions

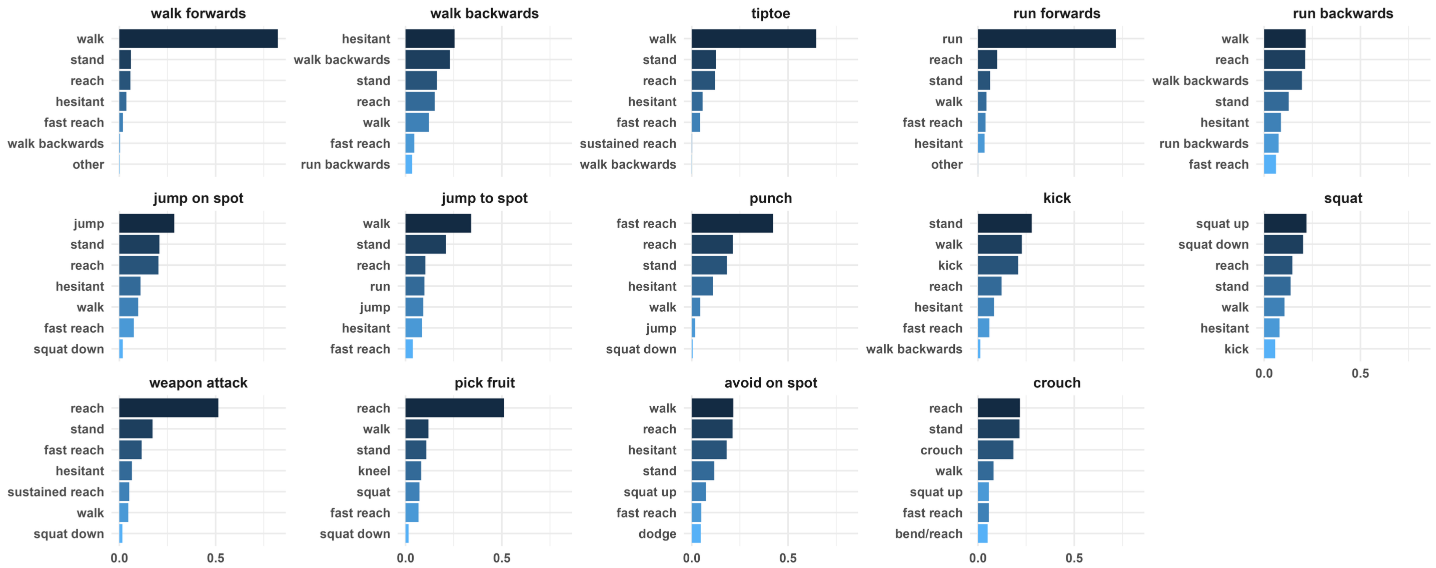


**Figure S1:** Instructed actions and the most frequent detected actions.

Figure S1 shows detected actions for the 14 instructed actions, averaged over participants and epochs. Instructions are listed *verbatim* in table S1. Notably, many instructions mandated more than one action such as walking to a particular spot to pick fruit. Also, there was no time pressure, so participants could stand and orient themselves if they wanted. As such, the classifier was expected to detect several action categories per instructed epoch, rather than a single action label.

For 11/14 of the instructed actions, the most frequently performed actions aligned with the semantic content of the instructions, as explained in the following. Of the remaining 3 instructions, one (*avoid on spot*) did not mandate a unique action, and two resulted in more complex patterns of detected actions.

##### Actions directly corresponding to instructions (5)

For 5 of the instructed actions (*walk forward, run forward, punch, squat, and jump on spot*), the dominant observed action corresponded directly to the instructed behavior. For the instruction *punch*, the most frequent action was labelled *fast reach*, which is kinematically similar. Squatting was separated by the classifier into two directional actions: *squat up* and *squat down*.

##### Actions consistent with the instructions (3)

For 3 further instructions, the dominant action was consistent with the instructions. *Weapon attack* and *pick fruit* engendered actions classified as *reach*. *Tiptoeing* was not detected as a distinct action category in the classifier but was classified as *walk,* as it primarily involved forward locomotion.

##### Actions consistent with the instruction to perform multiple actions (3)

For 3 further instructions, they explicitly involved several different actions. The instructions for *walk backwards* involved first going forward to a spot, standing there, and then backward. Here, *walking backwards* was the second most frequent action, preceded by *hesitant*. For *kick*, participants were instructed to stand and kick a sphere. Here, *kick* was the third most frequent action, preceded by *stand* and *walk*. For *crouch*, participants were instructed to *walk* to a tunnel, crouch through it, and then touch three spheres. Here, *crouch* was the third most frequently detected action, preceded by *reach* and *stand*.

#### Instructions instigating complex patterns of detected actions (2)

For *jump to spot*, the instructions required repeated vertical jumping, combined with horizontal displacement. In practice and verified from video observations, participants often covered distance using large, bounding strides that blended jumping with forward locomotion. As a result, these movements were frequently classified as *walking* or *running*, whereas discrete vertical jumps were detected less often. This appears to be not a failure of the action segmentation but a deviation of participants' behaviour from the intended actions.

For *run backwards*, the detected actions were variable and notably different from *walk backwards*. To begin with, the instructions mandated going forward first, which was detected as the most frequent action. Although both *backward walking* and *backward running* require increased balance control compared to forward locomotion, the two movements differ in their kinematic profiles. *Walking backwards* typically involves shorter, more controlled steps and lower momentum, which results in relatively stable and easily classifiable movement patterns. In contrast, *running backwards* introduces greater momentum and dynamic instability, often prompting participants to reduce speed or shorten stride length to maintain balance. As a result, the executed movement often falls below the velocity or stride thresholds that distinguish *running* from *walking* – a distinction biomechanically rooted in the absence of double stance phase. Consequently, the action classifier classified many instances of *running backwards* as *walking backwards*. In addition, participants frequently extended their arms forward more while moving backward to stabilize posture. These stabilizing movements may have increased the detection of *reaching* actions, further reducing the relative frequency of the *running* category. Although *running backwards* was present, it occurred at a lower frequency relative to these stabilizing and locomotor behaviors.

#### Instructions not mandating a specific action (1)

For *avoid on spot*, the instruction was to avoid spheres appearing in the air, without locomotion, that is, by leaning sideways or backwards. Here participants first had to *walk* to a certain spot and wait for the spheres to appear before they could perform the instructed action. We observe a high frequency of walk, reach (reflecting stabilizing movements), hesitant (reflecting standing with torso movements) and stand.
